# Highly Sequence-specific, Timing-controllable m^6^A Demethylation by Modulating RNA-binding Affinity of m^6^A Erasers

**DOI:** 10.1101/2024.06.24.600547

**Authors:** Kenko Otonari, Yuri Asami, Kosuke Ogata, Yasushi Ishihama, Shiroh Futaki, Miki Imanishi

## Abstract

In recent years, significant progress has been made in developing tools that consist of programmable RNA binding proteins and m^6^A-erasers to selectively demethylate m^6^A at specific sites. Especially, timing-controllable demethylation tools using stimulus-inducible dimerization show promise in understanding the impact of individual m^6^A modifications on dynamic physiological processes. However, a critical issue with these technologies is the possible off-target effects, wherein wild-type m^6^A-erasers constituting the tools may decrease methylation levels at sites other than the intended target m^6^A. In this study, we addressed this issue by reducing the intrinsic RNA-binding ability of m^6^A-erasers to prevent off-target effects and by fusing with PUF RNA-binding protein to provide recognition ability of arbitrary RNA sequences. Taking advantage of the reduced RNA-binding ability of the m^6^A-erasers themselves, we employed a rapamycin-based switching system in the linker connecting the PUF and m^6^A-eraser domains. The resulting m^6^A-erasers exhibited superior precision in demethylating targeted sites while minimizing off-target effects, presenting a novel approach for precise temporal control over m^6^A modification dynamics.

## INTRODUCTION

*N*^6^-methyladenosine (m^6^A) is one of the most abundant post-transcriptional modifications in eukaryotic RNA, and its physiological roles have been a research focus. M^6^A is installed in its consensus sequence (RRACH; R: A/G, H: A/C/U) by a methyltransferase complex (m^6^A-writers) and removed by demethylases (m^6^A-erasers) dynamically(1–6). It is recognized by m^6^A binding proteins (m^6^A-reader) and controls alternative splicing(7), mRNA stability(8, 9), and translation efficiency(10). Studies manipulating the expression of the m^6^A-erasers, fat mass and obesity-associated protein (FTO) and AlkB homolog 5 (ALKBH5), revealed their crucial roles in regulating various biological phenomena, including spermatogenesis(11), tumorigenesis(12, 13), adipogenesis(14, 15), and differentiation(16, 17) through their demethylation activities. However, the identification of the m^6^As responsible for the phenotypes has been difficult because overexpression or knockdown of m^6^A-erasers can alter the m^6^A modification levels of global transcripts. To address this difficulty, programmable m^6^A demethylation tools have been developed utilizing sequence-specific RNA binding proteins that guide FTO or ALKBH5 to the vicinity of target m^6^As(18–20). More recently, tools have been developed that allow the timing-control of programmable m^6^A demethylation by associating the RNA-binding proteins and m^6^A-demethylases in response to stimuli(21, 22). These tools paved the way to demethylate an individual m^6^A at specific timing for exploring the dynamic regulation seen in sequential or stepwise phenomena such as differentiation, somatic cell reprogramming, circadian clock and cell cycle. On the other hand, the possible increase in background demethylations by the overexpressed m^6^A-eraser was not considered, even though the global decrease in m^6^A levels by overexpressing m^6^A-erasers in living cells has been reported(5, 23). Therefore, the development of engineered m^6^A-erasers that specifically exert their demethylating activity for a specific sequence at a desired timing is highly needed.

m^6^A-erasers have intrinsic RNA-binding ability and this leads to the off-target binding of the eraser and non-specific m^6^A demethylation. Hence, one promising approach to suppress the off-target effects of sequence specific m^6^A-erasers is to reduce the intrinsic RNA-binding ability, thereby enabling sequence-selective action of the m^6^A-eraser through an RNA-binding protein segment capable of binding to the RNA sequence of interest. A similar approach has been reported by Liu and coworkers on an m^6^A-”writer” consisting of catalytically dead Cas13 (dCas13) for targeting the specific RNA sequences and METTL3:METTL14 methyltransferase complex with deletion of the zinc finger RNA-binding domain of METTL3 to weaken its own RNA-binding ability(24). In contrast to the METTL3:METTL14 methyltransferase complex, in which RNA-binding domains exist independent of the active site(25–28), the RNA-binding and active sites of m^6^A-erasers are closely related structurally(3, 4, 29). Although reducing the RNA-binding ability is an effective strategy to suppress the off-target effects, it is not a straightforward task to weaken the RNA-binding ability while maintaining catalytic activity of the m^6^A-erasers.

In this report, we aimed to create m^6^A-erasers to achieve highly sequence-specific m^6^A demethylation. Two m^6^A-erasers, FTO and ALKBH5, were employed as templates. Their RNA-binding abilities were modulated based on their structural data to suppress the non-specific binding to undesirable off-target RNA sequences, and we obtained “modulated” RNA-binding (mod) FTO and modALKBH5. For the recognition of specific RNA segments, they were fused with the RNA-binding protein PUF; we have previously demonstrated the feasibility of this protein to recognize arbitrary RNA sequences by arranging the repeated protein domains within PUF(18, 30). PUF is smaller than dCas13 and does not require guide RNA. This may allow for the flexible designing and easy preparation of m^6^A-erasers. Importantly, a switching system using FK-506 binding protein (FKBP) and FKBP-rapamycin-binding domain of FKBP12-rapamycin associated protein (FRB) was installed in the linker to connect the PUF domain and the mod-m^6^A-erasers, allowing for time-controllable, sequence-specific m^6^A demethylation (Scheme 1).

## MATERIALS AND METHODS

### Plasmid designing

The *E. coli* expression vector of mouse FTO (His6-mFTO-Strep/pET28b) and FTO-PUF have been described in our previous report(18). The expression vectors of mutants were synthesized by PCR using mutagenesis primers and subcloned into the above plasmids. For designing modALKBH5-PUF, the same method was used as FTO. For construction of the expression vectors of imodFP and imodAP, FRB- and FKBP-encoding DNA fragments were subcloned as fused by 2GSS linker in pET28b vector, resulted in His6-FTO-2GSS-FRB, ALKBH5-2GSS-FRB-Strep and His6-FKBP-2GSS-PUF-Strep in pET28b.

### Recombinant protein preparation

Protein expression and purification were performed basically based on our previous research(18). *E. coli* BL21 (DE3) cells were transformed with one of the above plasmids and grown on LB-agar plates containing 25 mg/L kanamycin. Protein expression was induced by adding 0.1 mM IPTG at logarithmic growth phase and incubated overnight at 18 ºC and 100 rpm. The cells were fractionated to soluble fraction by sonication and centrifugation, and the fraction was purified by the HisTrap FF (Cytiva, #17531901) followed by StrepTrap HP (Cytiva, #29048653) or StrepTrap XT (Cytiva, #29401317). FTO, FTO-PUF, ALKBH5-PUF and their mutants were concentrated with Amicon Ultra – 0.5 mL 30kDa (Millipore, #UFC503024) using 25 mM Tris-HCl (pH 7.5). FTO-FRB, modFTO-FRB, ALKBH5-FRB, modALKBH5-FRB and FKBP-PUF were concentrated using 25 mM Tris-HCl (pH 7.5) with 100 mM NaCl.

### RNA isolation

Total RNA was extracted from the cultured HEK293T cells using NucleoSpin RNA Plus (MACHEREY-NAGEL, #740984). mRNA was isolated using Dynabeads Oligo(dT)_25_ (Invitrogen, #61005) following the manufacture’s protocols. The concentrations of mRNA were measured by Qubit 4 Fluorometer (Invitrogen).

### *In vitro* demethylation and validation by MazF cleavage assay

50 nM of on-target RNA and off-target RNA (see Supplementary Table 1) were incubated with the indicated concentrations of demethylases in demethylation buffer (total 10 µL). The buffer composition for validating the demethylation activities of FTO-PUF and its mutants was 25 mM Tris-HCl (pH 7.5), 35 µM Fe(NH_4_)_2_(SO_4_)_2_·6H_2_O, 50 µM α-ketoglutarate, 500 µM L-ascorbate, 50 mM NaCl, 50 ng/µL BSA, 0.01% Tween20, 50 ng/µL total RNA from HeLa cells, and for ALKBH5-PUF and mutants, imodFP and imodAP was 25 mM Tris-HCl (pH 7.5), 283 µM Fe(NH_4_)_2_(SO_4_)_2_·6H_2_O, 300 µM α-ketoglutarate, 2 mM L-ascorbate, 50 mM NaCl, 50 ng/µL BSA, 0.01% Tween20, 50 ng/µL total RNA from HeLa cells, at 25 ºC for 1 hr followed by heating at 95 ºC for 3 min to stop reaction. For validating the demethylation activity of wild-type FTO or modFTO towards m^6^A on mRNA *in vitro*, the mRNA was demethylated in the following buffer: 25 mM Tris-HCl (pH 7.5), 283 µM Fe(NH_4_)_2_(SO_4_)_2_·6H_2_O, 300 µM α-ketoglutarate, 2 mM L-ascorbate, 50 mM NaCl, 0.5 U/µL RNasin Plus (Promega, #N261A) with 1 µM protein, at 37 ºC for 10 min, then reaction was quenched by adding 5 mM EDTA.

For MazF assays, 2 µL of the demethylated RNA was subjected to 8 µL of MazF buffer: 40 mM PBS (pH 7.5), 5 mM EDTA, 250 nM MazF, and incubated at 37 ºC for 1 hr followed by heating at 95 ºC for 3 min. The MazF-digested sample was mixed with the same amount of Hi-Di Formamide (Applied Biosynthesis, #4311320), heated at 95 ºC for 3 min, and immediately cooled on ice. 8 µL of the sample was loaded onto 20% urea-polyacrylamide gel and electrophoresed in 0.5 × TBE buffer (Nippon Gene, #318-90041). The fluorescently labeled RNAs were visualized using Amersham Typhoon (GE healthcare).

### Fluorescence polarization assay

20 nM of 5 ′ -FAM-labeled m^6^A-modified ssRNA was incubated with increasing concentrations (40 to 3×10^4^ nM) of wild-type FTO or modFTO in FP buffer (25 mM Tris-HCl, 0.01% Tween20, pH 7.5) for 1 hr at 25 ºC in a 96-well half-area microplate (Corning, #CLS3694). Fluorescence anisotropy was measured on an Infinite F Plex (TECAN) with a 485 nm excitation light and a 535 nm fluorescence wavelength filter. The results were analyzed by Kaleida graph (Synergy software, version 4.5.2.), and the binding dissociation constant was calculated by the following equation.

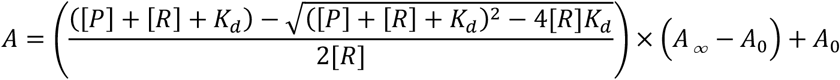

[P]: Protein concentration, [R]: FAM-labeled ssRNA concentration, A: Anisotropy

### Measurement of m^6^A level of mRNA using LC-MS/MS

The demethylated mRNA was purified using TRIzol LS Reagent (Invitrogen, #10296010) according to the manufacture’s protocol and the concentration was measured by Nanodrop. 400 ng of the purified mRNA was decapped with 20 units of RppH (NEB, #M0356) in 1× Thermopol buffer (TritonX-100 concentration was reduced to 0.01%) at 37 ºC for 6 hr. Decapped mRNA was digested to single nucleotides with 50 units Nuclease P1 (NEB, #M0660) in 20 mM NH_4_Ac at 37 ºC for 12 hr, and then 5’ phosphates of the nucleotide sample were removed with 0.5 units of quick-CIP (NEB, #M0525) at 37 ºC for 2 hr with agitation at 800 rpm for 1 min every 10 min.

Nucleoside sample was desalted by solid phase extraction (SPE) using Sep-Pak tC_18_ 1 cc Vac Cartridge (Waters, #WAT036820). The sample was diluted with SPE buffer A (0.1% heptafluorobutyric acid (HFBA) in H_2_O) to 1 mL. The cartridge was activated with 1-mL MeOH and SPE buffer B (0.1% HFBA, 80% acetonitrile (ACN)), and then equilibrated with 1-mL SPE buffer A. The diluted nucleoside was bound to the cartridge, washed with 1-mL SPE buffer A, and subsequently eluted by 1-mL SPE buffer B to 1.5-mL tube. The purified sample was evaporated using SpeedVac and dissolved in 20-µL mobile phase A and 10 µL was injected into an LC-MS/MS system consisting of dual LC-30AD pumps, a SIL-30AC autosampler, a CTO-30A column oven, an SPD-M20A photodiode array detector and an LCMS-8060 triple quadrupole mass spectrometer (SHIMADZU). The mobile phases consisting of 0.1% formic acid in H_2_O (A) and in ACN (B) were used. The sample was separated on an L-column 3 C18 (2.1 mm ID, 150 mm length, 2 µm C18-silica, 12 nm pore) (CERI) with a gradient program of 0-10 % B over 15 minutes. The flow rate was 200 µL/min. The mass spectrometer was operated in multiple reaction monitoring (MRM) mode with a dwell time of 100 ms and collision energy of -40. Adenosine and m^6^A were quantified with the transitions 268.1033 -> 136.0614 and 282.1191 -> 150.0773, respectively.

## RESULTS AND DISCUSSION

### Creation of modulated-RNA-binding (mod) FTO-PUF

FTO interacts with RNA at a positively charged groove-like region, and the crystal structure of FTO shows that the basic residues (K88, K213) at the groove-like region form the ‘pincer structure’, which can hold substrate RNA strands(6, 29). To reduce the RNA-binding affinity of FTO itself, alanine substitutions were introduced into FTO at the basic residues (K88, K213) in the ‘pincer structure’ (Figure 1A). The FTO mutants were fused with the sequence-specific RNA binding protein, PUF, binding to the UGUAUAUA sequence, as we reported previously(18). The demethylation activity of PUF-fused FTO mutants was examined toward the on-target m^6^A close to the PUF-binding site and the off-target m^6^A lacking a PUF-binding site, using the m^6^A-sensitive endoribonuclease MazF(31) (Figure S1A, Table S1). Wild-type FTO-PUF (FP) showed sequence-specific demethylation at lower than 500 nM but remarkable off-target demethylation occurred at higher concentrations (Figure 1B, C, Figure S1B). FTO^K88A^-PUF also demethylated both on- and off-target m^6^As at high concentrations, similar to FP (Figure 1B, Figure S1B, C), whereas FTO^K213A^-PUF showed a slightly reduced off-target effect while retaining its demethylation activity toward the on-target RNA (Figure 1B, Figure S1B, D). Interestingly, the combination of these alanine substitutions, FTO^K88A/K213A^-PUF, successfully reduced off-target demethylation activity even at high concentrations while maintaining on-target m^6^A demethylation activity (Figure 1B, C). K88 and K213 may cooperatively contribute to interactions with the substrate RNA. It is plausible that FTO^K88A/K213A^-PUF relies on PUF for RNA binding and that FTO^K88A/K213A^ exhibits high sequence selectivity by guided to its target m^6^A via PUF. We named the alanine substitution, FTO^K88A/K213A^, as ‘modified-RNA-binding FTO (modFTO)’ (Figure 1A, D).

**Figure 1.**
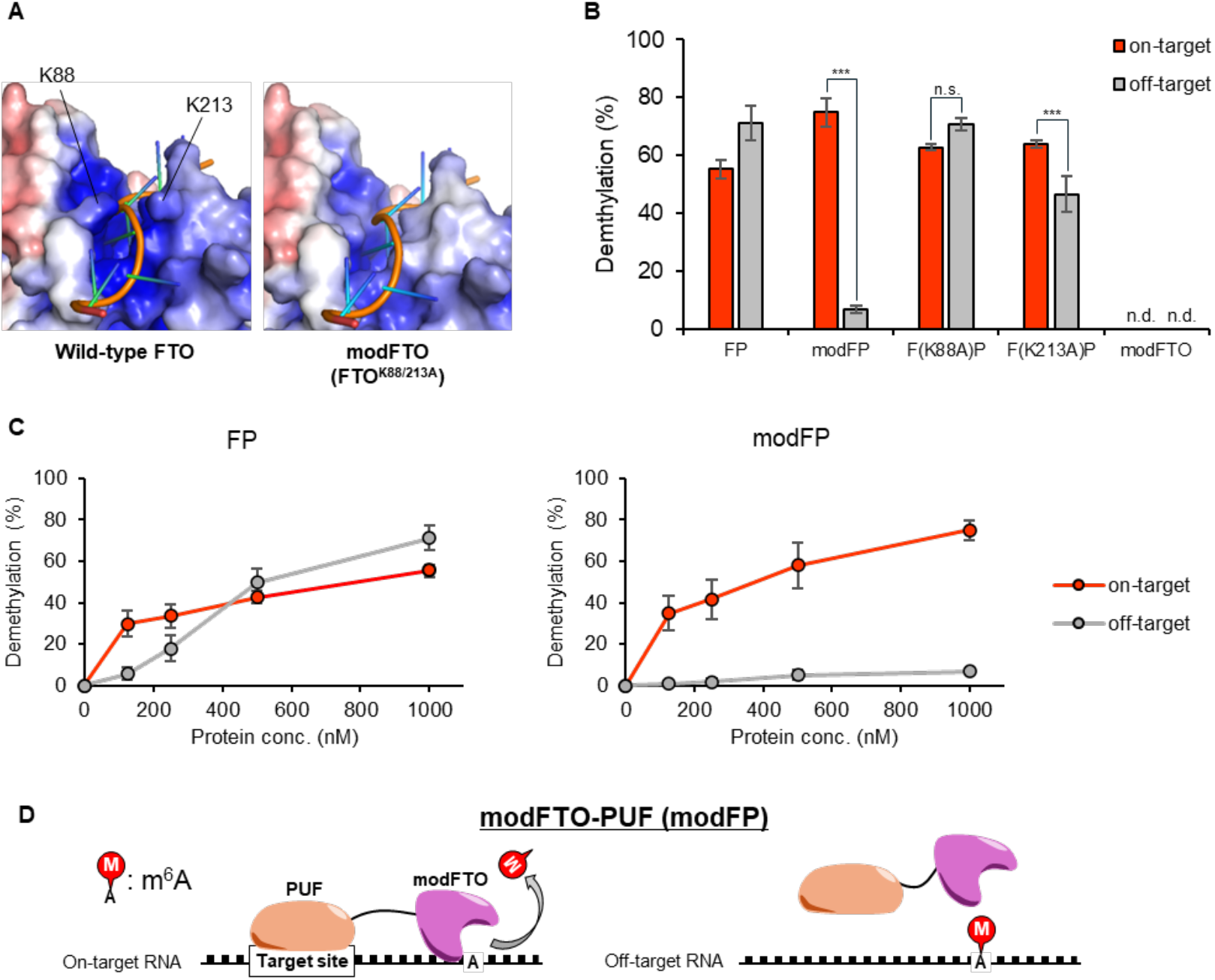
Creation of modFTO based on RNA recognition residues. (**A**) The pincer structure of wild-type FTO consisting of the two lysine residues (K88 and K213) interacting with the phosphate backbone of the ssRNA (left; PDB ID: 3LFM) and its corresponding region of modFTO (right; modified from 3LFM). (**B, C**) The sequence-specific demethylation activities to on-target RNA or off-target RNA of the alanine substitutions. (B) 1 mM or (C) indicated concentrations of FTO-PUF (FP) or modFP were incubated with two m^6^A-modified RNA oligos at 25ºC for 1 h. n.d. stands for not detected. Values and error bars indicate mean ± SEM (n = 4, Tukey; ***: p < 0.001). (**D**) Schematic illustration about the sequence-specific m^6^A demethylation by modFP.

### Characterization of modFTO

The two alanine substitutions were introduced to reduce the RNA-binding affinity, and then we measured the *K*_d_ values of wild-type FTO and modFTO for ssRNA by fluorescence polarization assay with FAM-labeled ssRNA (Table S1). Wild-type FTO and modFTO showed 0.93 ± 0.39 *μ*M and 2.89 ± 0.64 *μ*M, respectively (Figure 2).

**Figure 2.**
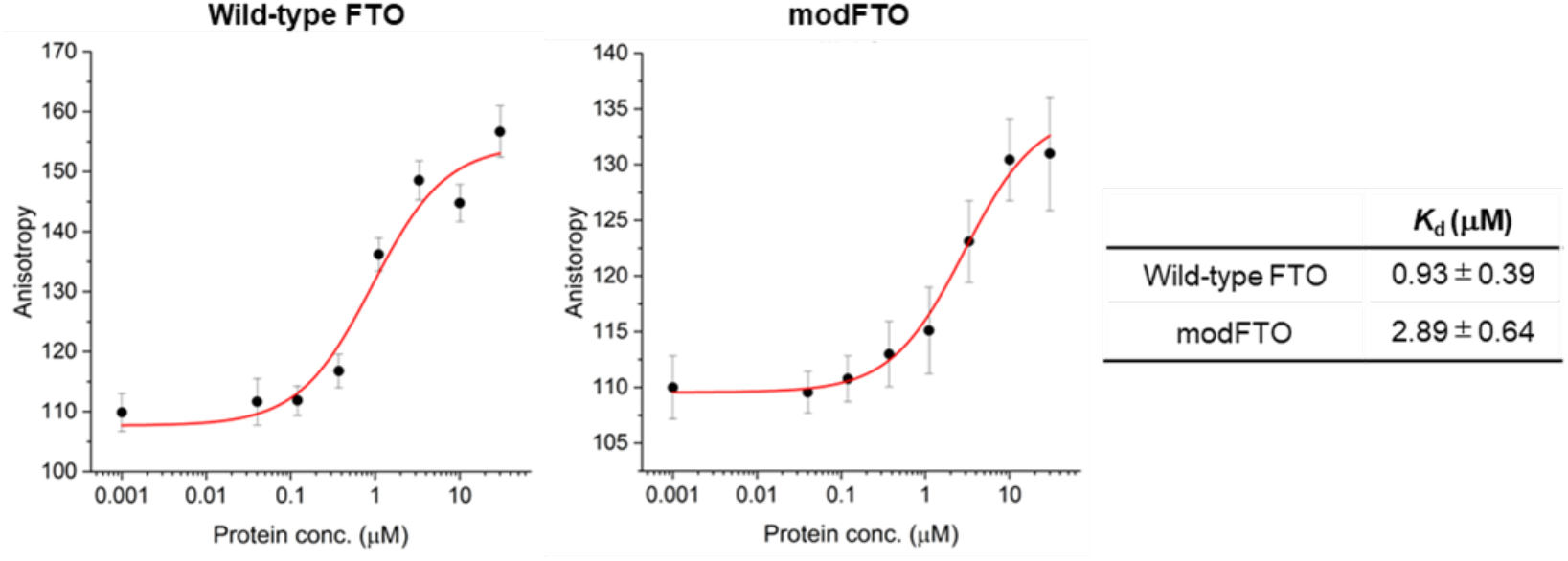
RNA-binding affinities of wild-type FTO and modFTO by fluorescence polarization assay. The binding affinity was measured as a change in anisotropy. Values and error bars indicate mean ± SEM (n = 3).

ModFTO itself, without PUF, showed little demethylation activity towards both on- and off-target m^6^As, even at high concentrations (Figure S2). Furthermore, we validated the demethylation activity towards mRNA transcribed in cells, not the synthesized oligos. mRNA isolated from HEK293T cells was demethylated and the ratios of m^6^A to A (m^6^A/A) was measured using liquid chromatography tandem mass spectrometry (LC-MS/MS). Wild-type FTO demethylated approximately 70% of m^6^A in mRNA, whereas modFTO did not show any significant demethylation (Figure S3A, B). These results suggest that the reduction in the RNA-binding affinity of modFTO compared to wild-type FTO is a possible reason for the increased specificity of modFP towards m^6^A near the PUF binding site and modFTO has lower background demethylation activity than wild-type FTO toward miscellaneous m^6^A derived from cells. In previous reports, overexpression or knockdown of FTO altered the m^6^A levels of global transcripts(5, 23). This lower background activity of modFTO toward miscellaneous m^6^A may contribute to the reduced off-target effects of modFP, even in living cells.

### Creation of modALKBH5

The generality of this design strategy for highly sequence-specific demethylation was verified using another m^6^A-eraser, ALKBH5. As expected, the fusion of wild-type ALKBH5 with the PUF RNA-binding protein, ALKBH5-PUF (AP), showed sequence-specific demethylation below 500 nM but at higher concentrations, it showed off-target effects owing to its innate recognition ability, similar to FP (Figure 3A, B). Therefore, we aimed to create a modulated RNA-binding ALKBH5 (modALKBH5) for higher sequence-specific m^6^A demethylation. Although both ALKBH5 and FTO belong to the AlkB subfamily, the pincer structure that was mutated in modFTO is unique to FTO among AlkB subfamily members(29) (Figure S4). In addition, the direction of the RNA phosphoribosyl backbone is reversed in ALKBH5 compared to that of the substrate structure of FTO(4). Therefore, we explored the unique residues that contribute to RNA binding in ALKBH5.

**Figure 3.**
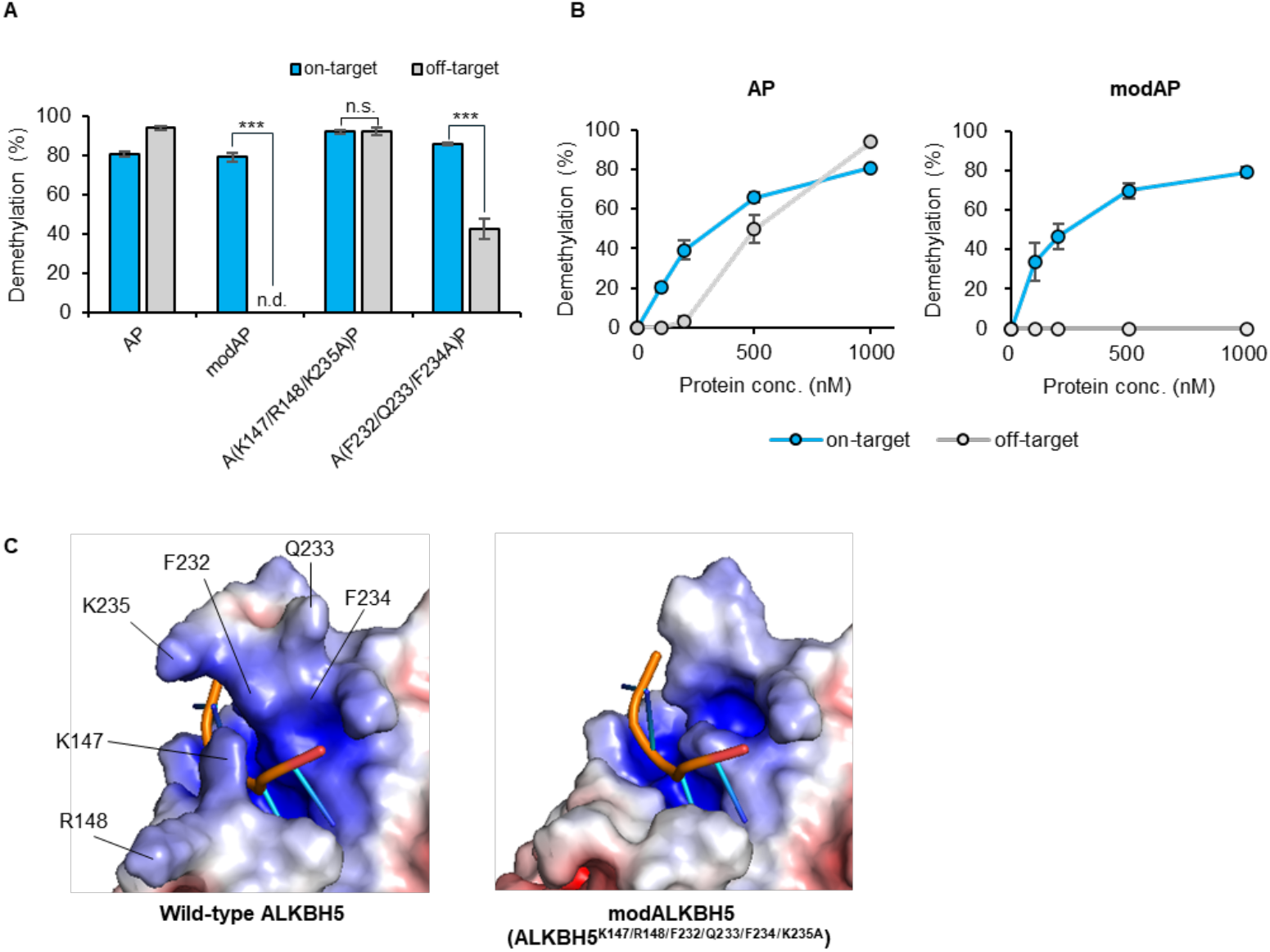
Creation of modAP based on RNA recognition domains. (A, B) The demethylation activities of the alanine-substitutions to on-target RNA or off-target RNA. (A) 1 *μ*M or (B) indicated concentrations of AP, modAP (= A(K147/R148/F232/Q233/F234/K235A)P) or each substitutions were incubated with two m^6^A-modified RNA oligos at 25ºC for 1 h. (C) The electrostatic surface of wild-type ALKBH5 (PDB ID; 7WL0) and modALKBH5 (modified from 7WL0). The six amino acids substituted with alanine in creating modALKBH5 are indicated. Values and error bars indicate mean ± SEM (n = 4, Tukey; n.s.: not significant, ***: p < 0.001).

Recent analysis of the crystal structure of ALKBH5 in complex with an m^6^A-containing ssRNA by Kaur *et al*. (4) suggested that nucleotide recognition lids (NRL) 2 and βIV-V loop are involved in the recognition of substrate ssRNA through electrostatic and π-π Stacking interactions (Figure 3C). Among the PUF-fused ALKBH5 mutants with alanine substitutions in the NRL2 and β IV - V loop regions, a mutant with six substitutions, ALKBH5^K147A/R148A/F232A/Q233A/F234A/K235A^-PUF, overcame the off-target effect (Figure 3A, B, Figure S5A), demethylating the on-target m^6^A but not the off-target m^6^A even at 1 *μ*M. We named this substituent as modALKBH5. Of the six substituted residues, the partial substitutions at K147/R148/K235, which contribute to the electrostatic interaction with RNA, still showed demethylation activity against off-target RNA at a concentration of 1 *μ*M (Figure 3A, S5A, B). This result is partly in accordance with a previous report demonstrating that ALKBH5^K147A^ did not reduce its demethylation activity compared to wild-type ALKBH5(3). ALKBH5^F232A/Q233A/F234A^-PUF, which has mutations at the π-π stacking interaction sites of ALKBH5, largely reduced the off-target effect (∼40%) while retaining the demethylation activity against on-target RNA (Figure 3A, Figure S5A, B). Our results showed that modALKBH5, in which all six residues were substituted with alanine, demethylated a small amount of m^6^A on its own (Figure S2), whereas modALKBH5-PUF (modAP) achieved higher sequence-selective demethylation that was dependent on the PUF binding site. This suggests that ALKBH5 recognizes its substrate RNAs through multiple residues, including not only basic amino acids but also aromatic ones. Furthermore, the results for modAP, as well as modFP, demonstrated the effectiveness of the strategy to reduce the RNA-binding ability of the m^6^A-erasers for highly sequence-specific m^6^A demethylation with low off-target effects.

### Rapamycin-inducible Sequence-specific m^6^A Eraser

Ligand-responsive m^6^A demethylation of a specific sequence is useful for switching the methylation state of target sites at the desired time. A rapamycin-inducible modFTO-PUF (imodFP) was designed using modFTO, which does not actively demethylate m^6^A by itself owing to its reduced RNA-binding ability (Figure S2). FK-506 binding protein (FKBP) and FKBP-rapamycin-binding domain of FKBP12-rapamycin associated protein (FRB) are known to dimerize rapidly in the presence of rapamycin(32). In imodFP, modFTO and PUF were fused to FRB and FKBP, resulting in modFTO-FRB and FKBP-PUF, respectively (Scheme 1). Evaluation of demethylation levels using the MazF assay showed that imodFP (modFTO-FRB and FKBP-PUF) successfully demethylated m^6^A on on-target RNA in the presence of rapamycin but not off-target m^6^A (Figure 4A, S6A). In contrast, iFP (wild-type FTO-FRB and FKBP-PUF) demethylated both on- and off-target m^6^As, regardless of the presence of rapamycin (Figure S6A, C). This may be because the wild-type FTO fused with FRB can demethylate essentially any m^6^As.

**Figure 4.**
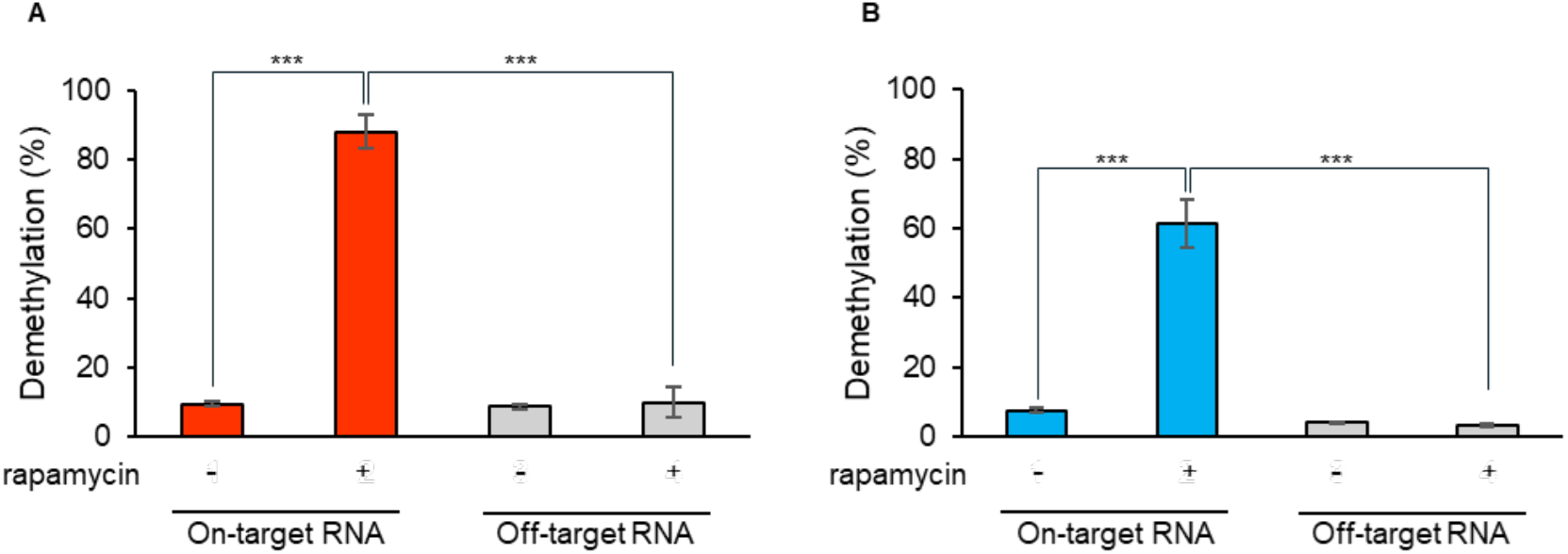
Rapamycin-inducible sequence-specific demethylation by imodFP. (A, B) Rapamycin-responsiveness of (A) imodFP or (B) imodAP. 1 *μ* M of modFTO-FRB or modALKBH5-FRB and FKBP-PUF were incubated with m^6^A-modified RNA oligos. Although imodFP and imodAP did not show demethylation activities in the absence of rapamycin, both showed activities to only on-target RNA in the presence of rapamycin. Values and error bars indicate means ± SEM (n=3, Turkey’s; ***: *p* < 0.001).

In the same strategy, we created imodAP (modALKBH5-FRB and FKBP-PUF) and examined its sequence-specificity and rapamycin-responsiveness. imodAP showed sequence-specific demethylation only in the presence of rapamycin (Figure 4B, S6B). On the other hand, iAP (wild-type ALKBH5-FRB and FKBP-PUF) did not show sequence-specificity or rapamycin-responsiveness as expected (Figure S6B, D). The high sequence- and ligand-dependent demethylation ability of imodFP and imodAP appears to rely on the properties of modFTO and modALKBH5, that do not demethylate on its own unless coupled with PUF (Figure S2). These results suggest that imodFP and imodAP can demethylate a specific m^6^A at arbitrary timing with high accuracy compared with other regulation tools.

## CONCLUSION

In summary, we created the regulation system of the timing of sequence-specific m^6^A demethylation with little off-target effect through engineering m^6^A-erasers whose RNA-binding affinity was finely tuned for the first time. To reduce the RNA-binding affinity, residues on the RNA recognition surface, but not in its active site, were substituted with alanine, so that they exert their demethylation ability only when in proximity to RNA. When in combination with PUF, the modulated m^6^A-erasers showed a highly efficient PUF-dependent m^6^A demethylation ability, with remarkable little off-target demethylation. To date, approaches to reduce non-specific substrate recognition of enzymes have been employed in the target RNA methylation (TRM) system using an m^6^A-writer lacking the zinc finger domain(24) and in highly target-specific genome-editing systems using Cas9 mutants with substituted residues in the substrate recognition or Rec3 domains(33–35). In contrast to these approaches, which use enzymes with independent substrate-recognition and catalytic domains, our strategy could be applied to other epitranscriptomic and epigenetic enzymes, especially when the catalytic sites and nucleic acid-recognition sites are closely related, while optimization is required for each enzyme.

Ultimately, the ligand-inducible imodFP and imodAP were designed by utilizing modFTO and modALKBH5. The imodFP and imodAP were activated by rapamycin treatment and showed highly sequence-specific demethylation. Targeted m^6^A demethylation systems in response to stimuli, such as abscisic acid and light irradiation, have been reported but these systems utilize wild-type m^6^A-erasers and do not consider background demethylation(21, 22). Our mod-m^6^A-erasers with reduced RNA-binding ability are expected to achieve more accurate timing- and sequence-specific demethylation of m^6^A than wild-type m^6^A-erasers in combination with various stimulus-responsive systems. Reprogramming of somatic cells into induced pluripotent stem cells is regulated by m^6^A modifications at specific periods(16, 17). Given its high selectivity for the target m^6^A and its ability to control demethylation timing, imodFP and imodAP would be a powerful tool to reveal the period-dependent roles of individual m^6^A modifications in dynamic physiological processes.

---

**Scheme 1.**
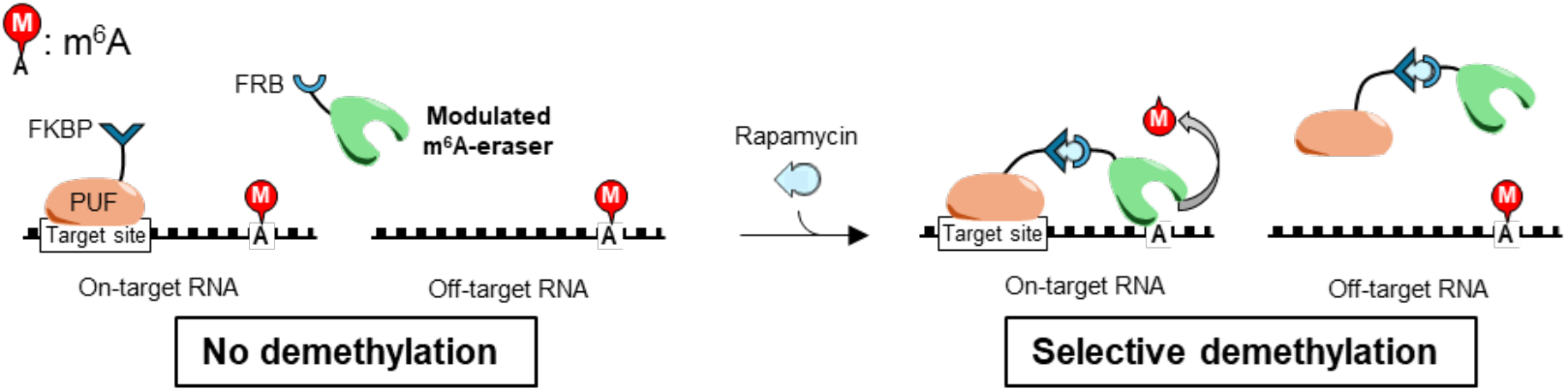
Concept of ligand-inducible sequence-specific m^6^A demethylation tool with minimal off-target effect. Modulated m^6^A-erasers do not demethylate in the absence of ligands but only demethylate the m^6^A nearby the PUF-binding site in the presence of ligands.

## REFERENCES

1. Dominissini, D., Moshitch-Moshkovitz, S., Schwartz, S., Salmon-Divon, M., Ungar, L., Osenberg, S., Cesarkas, K., Jacob-Hirsch, J., Amariglio, N., Kupiec, M., et al. (2012) Topology of the human and mouse m6A RNA methylomes revealed by m6A-seq. Nature, 485, 201–206.

2. Liu, J., Yue, Y., Han, D., Wang, X., Fu, Y., Zhang, L., Jia, G., Yu, M., Lu, Z., Deng, X., et al. (2014) A METTL3–METTL14 complex mediates mammalian nuclear RNA N6-adenosine methylation. Nat. Chem. Biol., 10, 93–95.

3. Feng, C., Liu, Y., Wang, G., Deng, Z., Zhang, Q., Wu, W., Tong, Y., Cheng, C. and Chen, Z. (2014) Crystal structures of the human RNA demethylase Alkbh5 reveal basis for substrate recognition. J. Biol. Chem., 289, 11571–11583.

4. Kaur, S., Tam, N. Y., McDonough, M. A., Schofield, C. J. and Aik, W. S. (2022) Mechanisms of substrate recognition and N6-methyladenosine demethylation revealed by crystal structures of ALKBH5–RNA complexes. Nucleic Acids Res., 50, 4148–4160.

5. Jia, G., Fu, Y., Zhao, X., Dai, Q., Zheng, G., Yang, Y., Yi, C., Lindahl, T., Pan, T., Yang, Y. G., et al. (2011) N6-Methyladenosine in nuclear RNA is a major substrate of the obesity-associated FTO. Nat. Chem. Biol., 7, 885–887.

6. Han, Z., Niu, T., Chang, J., Lei, X., Zhao, M., Wang, Q., Cheng, W., Wang, J., Feng, Y. and Chai, J. (2010) Crystal structure of the FTO protein reveals basis for its substrate specificity. Nature, 464, 1205–1209.

7. Xiao, W., Adhikari, S., Dahal, U., Chen, Y.-S., Hao, Y.-J., Sun, B.-F., Sun, H.-Y., Li, A., Ping, X.-L., Lai, W.-Y., et al. (2016) Nuclear m6A reader YTHDC1 regulates mRNA splicing. Mol. Cell, 61, 507–519.

8. Wang, X., Lu, Z., Gomez, A., Hon, G.C., Yue, Y., Han, D., Fu, Y., Parisien, M., Dai, Q., Jia, G., et al. (2014) N6-methyladenosine-dependent regulation of messenger RNA stability. Nature, 505, 117–120.

9. Du, H., Zhao, Y., He, J., Zhang, Y., Xi, H., Liu, M., Ma, J. and Wu, L. (2016) YTHDF2 destabilizes m6A-containing RNA through direct recruitment of the CCR4–NOT deadenylase complex. Nat. Commun., 7, 12626.

10. Shi, H., Wang, X., Lu, Z., Zhao, B. S., Ma, H., Hsu, P. J., Liu, C. and He, C. (2017) YTHDF3 facilitates translation and decay of N6-methyladenosine-modified RNA. Cell Res., 27, 315–328.

11. Zheng, G., Dahl, J. A., Niu, Y., Fedorcsak, P., Huang, C.-M., Li, C. J., Vågbø, C. B., Shi, Y., Wang, W.-L., Song, S.-H., et al. (2013) ALKBH5 is a mammalian RNA demethylase that impacts RNA metabolism and mouse fertility. Mol. Cell, 49, 18–29.

12. Yang, S., Wei, J., Cui, Y.-H., Park, G., Shah, P., Deng, Y., Aplin, A. E., Lu, Z., Hwang, S., He, C., et al. (2019) m6A mRNA demethylase FTO regulates melanoma tumorigenicity and response to anti-PD-1 blockade. Nat. Commun., 10, 2782.

13. Shen, C., Sheng, Y., Zhu, A. C., Robinson, S., Jiang, X., Dong, L., Chen, H., Su, R., Yin, Z., Li, W., et al. (2020) RNA demethylase ALKBH5 selectively promotes tumorigenesis and cancer stem cell self-renewal in acute myeloid leukemia. Cell Stem Cell, 27, 64–80. e9.

14. Wu, W., Feng, J., Jiang, D., Zhou, X., Jiang, Q., Cai, M., Wang, X., Shan, T. and Wang, Y. (2017) AMPK regulates lipid accumulation in skeletal muscle cells through FTO-dependent demethylation of N6-methyladenosine. Sci. Rep., 7, 41606.

15. Zhao, X., Yang, Y., Sun, B.-F., Shi, Y., Yang, X., Xiao, W., Hao, Y.-J., Ping, X.-L., Chen, Y.-S., Wang, W.-J., et al. (2014) FTO-dependent demethylation of N6-methyladenosine regulates mRNA splicing and is required for adipogenesis. Cell Res., 24, 1403–1419.

16. Batista, P.J., Molinie, B., Wang, J., Qu, K., Zhang, J., Li, L., Bouley, D.M., Lujan, E., Haddad, B., Daneshvar, K., et al. (2014) m6A RNA modification controls cell fate transition in mammalian embryonic stem cells. Cell Stem Cell, 15, 707–719.

17. Geula, S., Moshitch-Moshkovitz, S., Dominissini, D., Mansour, A.A., Kol, N., Salmon-Divon, M., Hershkovitz, V., Peer, E., Mor, N., Manor, Y.S., et al. (2015) m6A mRNA methylation facilitates resolution of naïve pluripotency toward differentiation. Science, 347, 1002–1006.

18. Shinoda, K., Suda, A., Otonari, K., Futaki, S. and Imanishi, M. (2020) Programmable RNA methylation and demethylation using PUF RNA binding proteins. Chem. Commun., 56, 1365–1368.

19. Rau, K., Rösner, L. and Rentmeister, A. (2019) Sequence-specific m6A demethylation in RNA by FTO fused to RCas9. RNA, 25, 1311–1323.

20. Li, J., Chen, Z., Chen, F., Xie, G., Ling, Y., Peng, Y., Lin, Y., Luo, N., Chiang, C. M. and Wang, H. (2020) Targeted mRNA demethylation using an engineered dCas13b-ALKBH5 fusion protein. Nucleic Acids Res., 48, 5684–5694.

21. Zhao, J., Li, B., Ma, J., Jin, W. and Ma, X. (2020) Photoactivatable RNA N6-methyladenosine editing with CRISPR-Cas13. Small, 16.

22. Shi, H., Xu, Y., Tian, N., Yang, M. and Liang, F.-S. (2022) Inducible and reversible RNA N6-methyladenosine editing. Nat. Commun., 13, 1–10.

23. Wei, J., Liu, F., Lu, Z., Fei, Q., Ai, Y., He, P. C., Shi, H., Cui, X., Su, R., Klungland, A., et al. (2018) Differential m6A, m6Am, and m1A demethylation mediated by FTO in the cell nucleus and cytoplasm. Mol. Cell, 71, 973–985.e5.

24. Wilson, C., Chen, P. J., Miao, Z. and Liu, D. R. (2020) Programmable m6A modification of cellular RNAs with a Cas13-directed methyltransferase. Nat. Biotech., 38, 1431–1440.

25. Wang, X., Feng, J., Xue, Y., Guan, Z., Zhang, D., Liu, Z., Gong, Z., Wang, Q., Huang, J., Tang, C., et al. (2016) Structural basis of N6-adenosine methylation by the METTL3– METTL14 complex. Nature, 534, 575–578.

26. Wang, P., Doxtader, K. A. and Nam, Y. (2016) Structural basis for cooperative function of Mettl3 and Mettl14 methyltransferases. Mol. Cell, 63, 306–317.

27. Śledź, P. and Jinek, M. (2016) Structural insights into the molecular mechanism of the m6A writer complex. Elife, 5, e18434.

28. Yoshida, A., Oyoshi, T., Suda, A., Futaki, S. and Imanishi, M. (2022) Recognition of G-quadruplex RNA by a crucial RNA methyltransferase component, METTL14. Nucleic Acids Res., 50, 449–457.

29. Zhang, X., Wei, L.-H., Wang, Y., Xiao, Y., Liu, J., Zhang, W., Yan, N., Amu, G., Tang, X., Zhang, L., et al. (2019) Structural insights into FTO’s catalytic mechanism for the demethylation of multiple RNA substrates. Proc. Nat. Acad. Sci. U.S.A., 116, 2919–2924.

30. Shinoda, K., Tsuji, S., Futaki, S. and Imanishi, M. (2018) Nested PUF proteins: extending target RNA elements for gene regulation. ChemBioChem, 19, 171–176.

31. Imanishi, M., Tsuji, S., Suda, A. and Futaki, S. (2017) Detection of N6-methyladenosine based on the methyl-sensitivity of MazF RNA endonuclease. Chem. Commun., 53, 12930–12933.

32. Banaszynski, L. A., Liu, C. W. and Wandless, T. J. (2005) Characterization of the FKBP-rapamycin-FRB ternary complex. J. Am. Chem. Soc., 127, 4715–4721.

33. Slaymaker, I. M., Gao, L., Zetsche, B., Scott, D. A., Yan, W. X. and Zhang, F. (2016) Rationally engineered Cas9 nucleases with improved specificity. Science, 351, 84–88.

34. Kleinstiver, B. P., Pattanayak, V., Prew, M. S., Tsai, S. Q., Nguyen, N. T., Zheng, Z. and Joung, J. K. (2016) High-fidelity CRISPR–Cas9 nucleases with no detectable genome-wide off-target effects. Nature, 529, 490–495.

35. Chen, J. S., Dagdas, Y. S., Kleinstiver, B. P., Welch, M. M., Sousa, A. A., Harrington, L. B., Sternberg, S. H., Joung, J. K., Yildiz, A. and Doudna, J. A. (2017) Enhanced proofreading governs CRISPR–Cas9 targeting accuracy. Nature, 550, 407–410.

