## Supplementary Figures for "Highly Sequence-specific, Timing-controllable m^6^A Demethylation by Modulating RNA-binding Affinity of m^6^A Erasers"

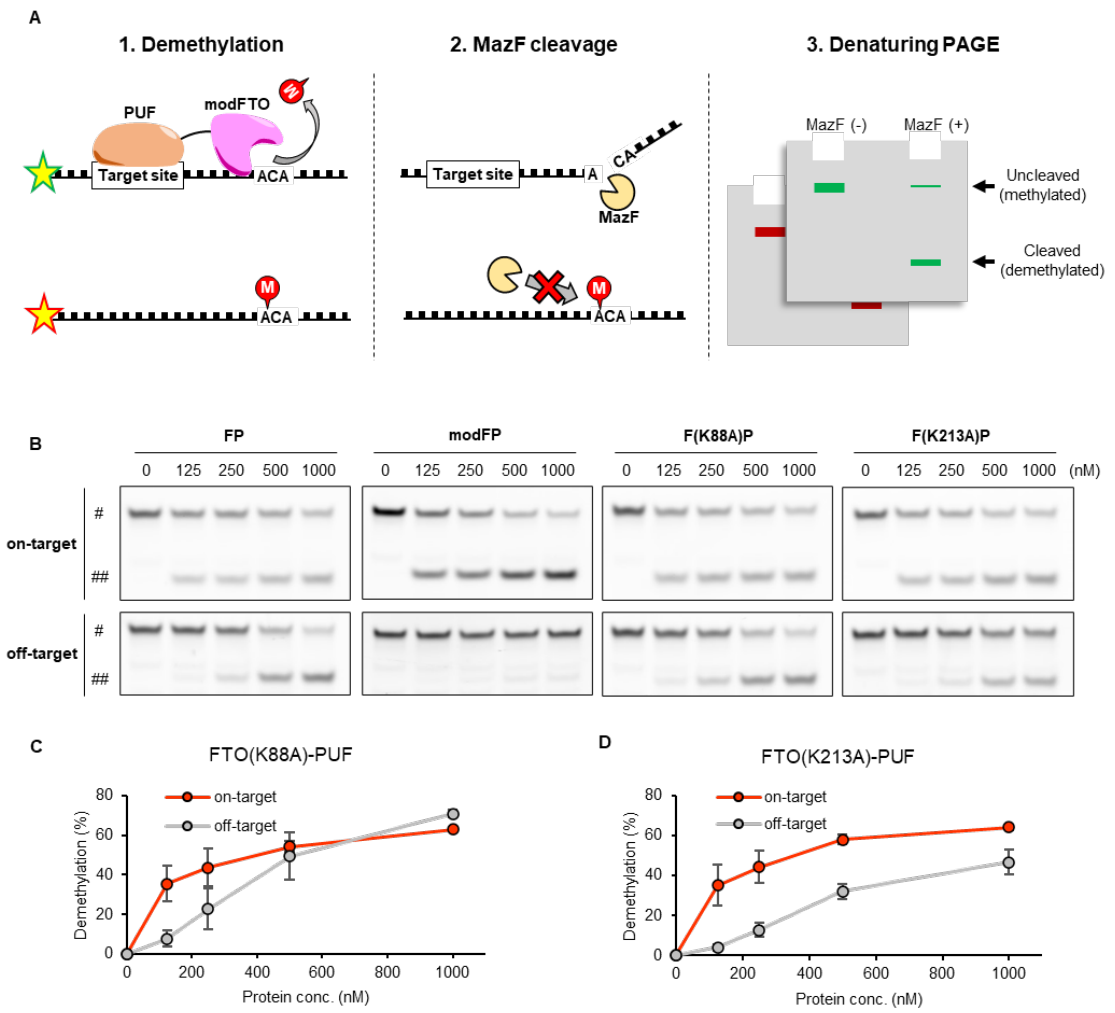


**Figure S1.** The demethylation activities of alanine substitutions to each m$^6$A-modified RNA oligos at each concentration. (**A**) Scheme of MazF assay for sequence-specific demethylation. MazF is an m6A sensitive RNA endonuclease and only cleaves demethylated single-strand RNA at ACA sequences. (**B**) PAGE images of MazF assay. On-target RNA and off-target RNA were visualized by excitation with 488 nm (upper) and 532 nm (lower). #: un-cleaved RNA (methylated RNA), ##: cleaved RNA (demethylated RNA). (**C, D**) The plot of concentration-dependent demethylation activities of (B) FTO(K88A)-PUF and (C) FTO(K213A)-PUF. Values and error bars indicate mean ± SEM (n=3).

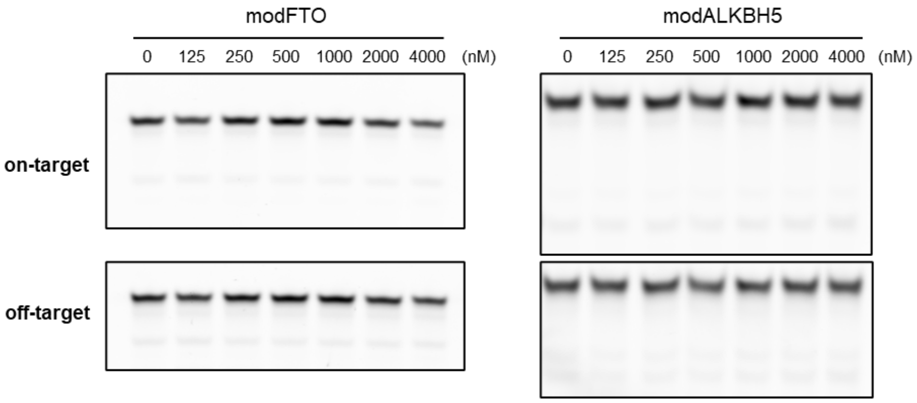


**Figure S2.** Demethylation assay of modFTO and modALKBH5 towards on-target RNA or off-target RNA evaluated by MazF digestion and PAGE analysis. Both $m^6A$-erasers did not show any demethylation activity to the two $m^6A$-modified RNA oligos even at high concentration (~ 4000 nM).

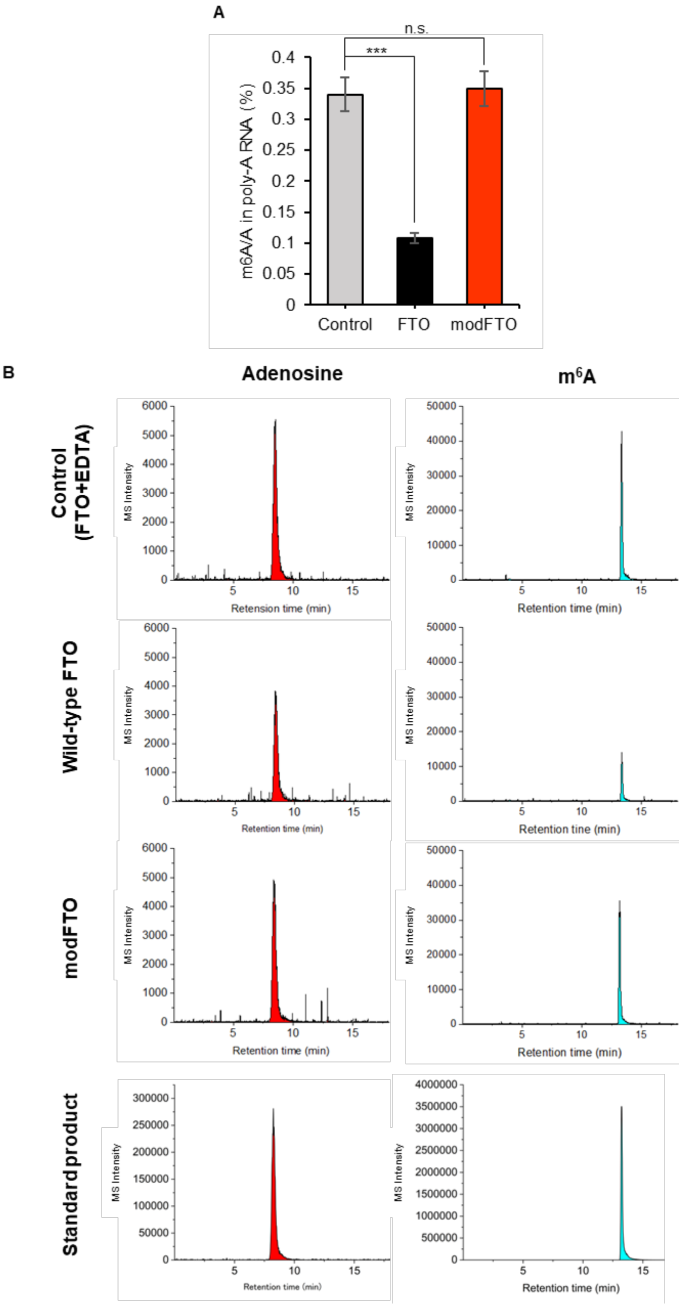


**Figure S3.** (A) Quantification of the m$^6$A/A in poly-A RNA by LC-MS/MS. While wild-type FTO showed significant decrease in m$^6$A/A compared to control (wild-type FTO + EDTA), and modFTO did not show any decrease. Values and error bars indicate mean ± SEM (n = 3, Turkey; n.s.: not significant, ***: p < 0.001). (B) Representative LC-MS/MS ion chromatogram of standard products (adenosine, m$^6$A) and mRNA purified from HEK293T cells treated with wild-type FTO or modFTO *in vitro*. Standard products were diluted at 1 g/L and injected 10 mL. The data were analyzed by OriginPro 2024 (ver. 10.1.0.170).

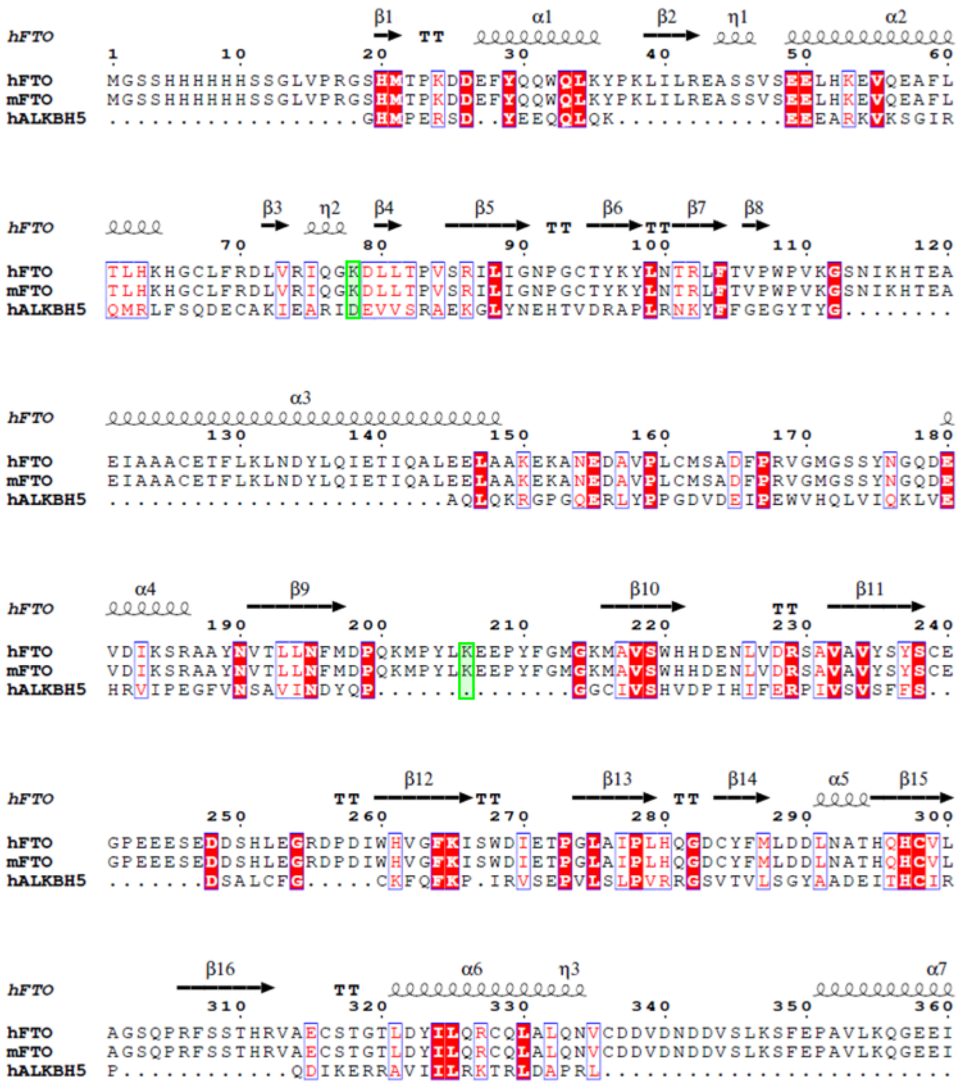

**Figure S4.** Structural-based sequence alignment of m$^6$A-erasers by STRAP. Alanine-substituted residues added for generating modFTO are colored in light-green. The secondary structure obtained from PDB file (ID: 3LFM) is displayed using ESPript 3.0[1]. hFTO: human FTO, mFTO: mouse FTO, hALKBH5: human ALKBH5.

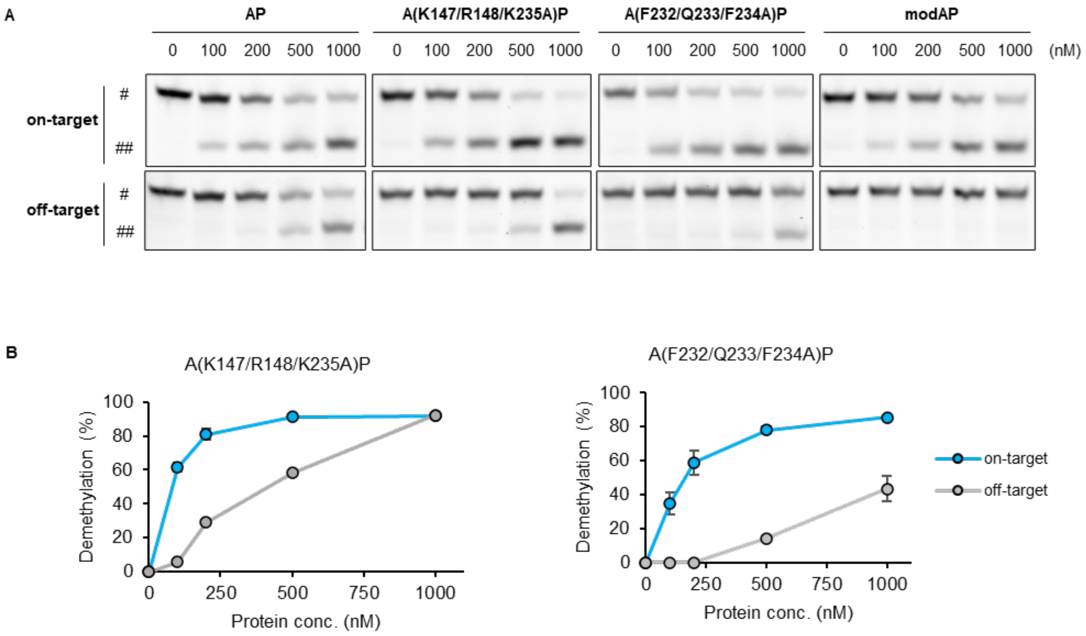


**Figure S5.** The demethylation activities of AP and its alanine substitutions to each m$^6$A-modified RNA oligos. (A) PAGE images of MazF assay. On-target RNA and off-target RNA were visualized by excitation with 488 nm (upper) and 532 nm (lower). #: un-cleaved RNA (methylated RNA), ##: cleaved RNA (demethylated RNA). (B) Plots of the concentration-dependent demethylation activities of A(K147/R148/K235A)P and A(F232/Q232/F234A)P. Values and error bars indicate mean ± SEM (n = 3).

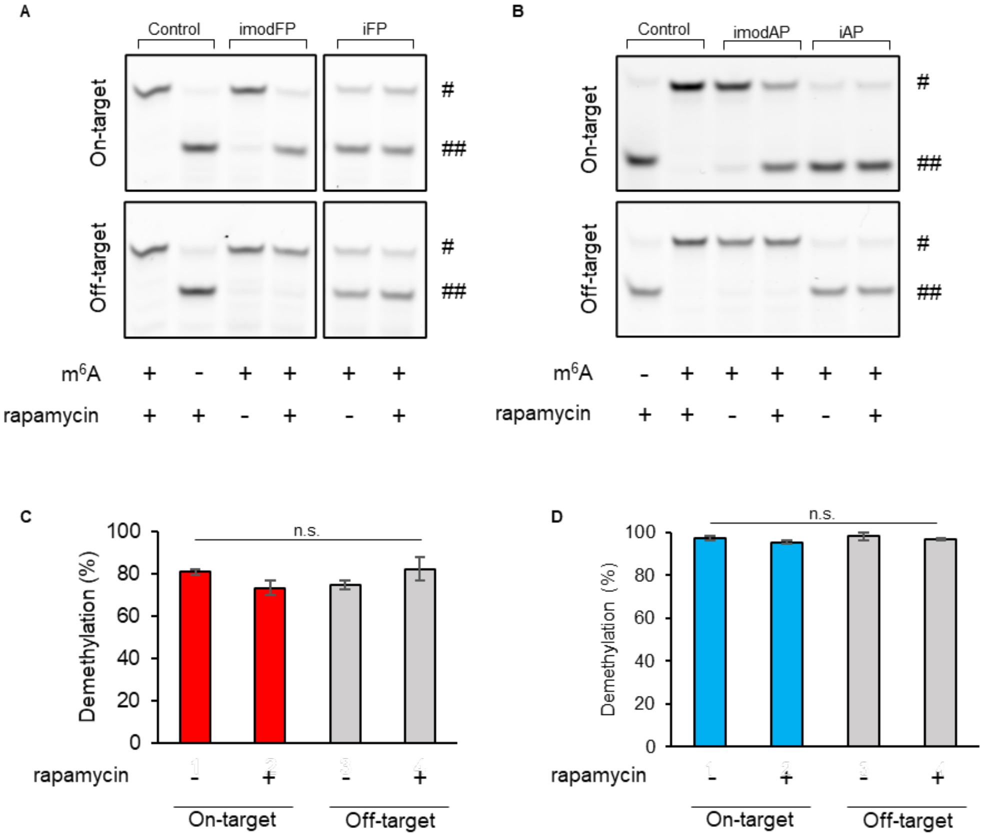


**Figure S6.** Sequence-specific demethylation activities and responsiveness to rapamycin of iFP, imodFP, iAP and imodAP. (A, B) PAGE images of MazF assay. The concentrations of (A) iFP and imodFP or (B) iAP and imodAP were 1 μM and control contained no protein. On-target RNA and off-target RNA were visualized by excitation with 488 nm (upper) and 532 nm (lower). #: un-cleaved RNA (methylated RNA), ##: cleaved RNA (demethylated RNA). (C, D) Demethylation efficiencies of (C) iFP and (D) iAP towards on-target RNA or off-target RNA. For validation of iAP's demethylation activity 2-nt RNA (Table S1) was used as on-target RNA. Values and error bars indicate mean ± SEM (n = 3, Turkey's; n.s.: not significant).

**Table S1** ssRNA oligo used for MazF assays and FP assay

| Oligo name | Sequence (5′ →3′ ) |
|---|---|
| on-target RNA | FAM-AU**UGUAUAUA**UCUAAG(m$^6$A)CAUUUUA |
| off-target RNA | TAMRA-AUAUCUCUUGGGUUCUAUUAG(m$^6$A)CAUUUAG |
| on-target MazF control | FAM-AU**UGUAUAUA**UCUAAGACAUUUUA |
| off-target MazF control | TAMRA-AUAUCUCUUGGGUUCUAUUAGACAUUUAG |
| on-target RNA 2 | FAM-AU**UGUAUAUA**AG(m$^6$A)CAUUUUA |
| RNA for FP assay | FAM-AUUGUAUAU(m$^6$A)CAUUUA |

UGUAUAUA: PUF binding sequence

### Reference


[1] Robert, X., and Gouet, P. (2014) Deciphering key features in protein structures with the new ENDscript server. *Nucleic Acids Res.*, **42**, 320–324.
